# Gsx2 Regulates Oligodendrocyte Precursor Formation in the Zebrafish Spinal Cord

**DOI:** 10.1101/2025.10.18.683250

**Authors:** Kimberly A. Arena, Christina A. Kearns, Mohamoud Ahmed, Rebecca O’Rourke, Charles G. Sagerström, Santos J. Franco, Bruce Appel

## Abstract

Nervous system development relies on sequential and coordinated formation of diverse neurons and glia from neural progenitor cells (NPCs). In the spinal cord, NPCs of the pMN domain produce neurons early in development followed by oligodendrocyte precursor cells (OPCs), which subsequently differentiate as oligodendrocytes (OLs), the myelinating glia of the central nervous system, later in development. The mechanisms that specify neural progenitor cells to the OL lineage are not yet well understood. Using zebrafish as an experimental model system, we generated single-cell RNA sequencing and single-nuclei ATAC sequencing data that identified a subpopulation of NPCs, called pre-OPCs, that appeared fated to produce OPCs. pre-OPCs uniquely express several genes that encode transcription factors specific to the OL lineage, including Gsx2, which regulates OPC formation in the mouse forebrain. To investigate Gsx2 function in zebrafish OPC specification, we used CRISPR/Cas9 genome editing to create *gsx2* loss-of-function alleles. *gsx2* homozygous mutant embryos initiated OPC formation prematurely and produced excess OPCs without altering OL differentiation. Using our single-nuclei multi-omics dataset, we predicted a gene regulatory network centered around *gsx2* and identified genes that might be transcriptionally regulated by Gsx2. Taken together, our studies suggest that Gsx2 expression in pre-OPCs contributes to the timing of OPC specification.

## INTRODUCTION

In the central nervous system (CNS) of vertebrate animals, axons are wrapped by myelin, a specialized, lipid-rich membrane produced by glial cells called oligodendrocytes. Myelin increases the conduction speed of electrical impulses through axons and supports neuronal health; therefore, oligodendrocytes are essential to vertebrate nervous system function. Accordingly, understanding how functional neural circuits are created during vertebrate development requires knowing how oligodendrocytes are produced.

Although oligodendrocytes occupy the entire CNS, they arise only from specific subpopulations of neural progenitor cells (NPCs). In the spinal cord, a ventral domain of NPCs known as pMN progenitors, which express the transcription factor-encoding gene *Olig2*, gives rise to oligodendrocyte precursor cells (OPCs) (Lu et al., 2000; Masahira et al., 2006; Park et al., 2002; Takebayashi et al., 2000; Zhou et al., 2000). OPCs divide and migrate to become dispersed throughout the spinal cord; eventually, many OPCs exit the cell cycle and differentiate as myelinating oligodendrocytes whereas others persist into adulthood (Rowitch, 2004).

Importantly, pMN progenitors also produce motor neurons in addition to some ventral interneurons, a subset of astrocytes, and ependymal cells (Lu et al., 2002; Masahira et al., 2006; Park et al., 2004; Zhou and Anderson, 2002). By analyzing clones of cells stemming from individually labeled progenitors at neural plate stage (approximately 10 hours post fertilization, hpf) in zebrafish, we found that progenitors that were close to the midline rarely divided and produced early born motor neurons and ventral interneurons (Park et al., 2004). By contrast, progenitors that were slightly further from the midline divided many times and individual clones contained both late-born motor neurons and OPCs (Park et al., 2004). Using fate-mapping and timelapse microscopy, we subsequently learned that by 24 hpf, after neural tube formation, motor neurons and OPCs arise from distinct progenitors (Ravanelli and Appel, 2015). Thus, between neural plate and neural tube stages, pMN progenitors become fate restricted, producing only neurons or only OPCs.

To determine the molecular identities of fate-restricted pMN progenitors, we performed single cell RNA-sequencing (scRNA-seq) of EGFP^+^ cells isolated from *Tg(olig2:egfp)* embryos. This revealed a small population of cells, which we called pre-OPCs, that appeared to represent a transition state between NPCs and OPCs (Scott et al., 2021). Among the genes uniquely expressed by pre-OPCs was *gsx2*, which encodes an evolutionarily conserved homeodomain transcription factor. In mice, *Gsx2*^+^ cells of the lateral ganglionic eminence (LGE) of the forebrain gave rise to oligodendrocyte lineage cells (OLCs) (Kessaris et al., 2006). LGE neurogenesis was impaired in mouse embryos lacking *Gsx2* function (Corbin et al., 2000; Toresson et al., 2000; Waclaw et al., 2009), accompanied by premature and excess OPC formation (Chapman et al., 2018; Chapman et al., 2012). Conversely, forcing telencephalic progenitors to express *Gsx2* blocked OPC formation (Chapman et al., 2012). Altogether, these observations indicate that Gsx2 regulates the time at which OPCs are produced by LGE progenitors.

Finding that zebrafish spinal cord pre-OPCs differentially express *gsx2* raised the question of whether Gsx2 functions similarly in forebrain and spinal cord to control OPC specification. To answer this question, we used CRISPR-Cas9 gene editing to evaluate oligodendrocyte development in zebrafish larvae lacking *gsx2* function. We also interrogated scRNA-seq and single-nucleus ATAC-seq data to learn how Gsx2 might contribute to a gene regulatory network for OPC specification.

## RESULTS

### pre-OPCs are a subset of neural progenitor cells that express *gsx2*

In zebrafish, pMN progenitors produce motor neurons from approximately 10 to 30 hours post fertilization (hpf) (Myers et al., 1986) (Figure 1A). OPC specification, revealed by expression of *sox10*, the earliest known marker of OPCs, begins by approximately 40 hpf (Scott et al., 2021) followed by differentiation of myelinating OLs by 72 hpf (Buckley et al., 2010) (Figure 1A). In a prior fate mapping study we determined that motor neurons and OPCs arise from distinct subpopulations of pMN progenitors (Ravanelli and Appel, 2015). Subsequently, we performed single-cell RNA-sequencing (scRNA-seq) of pMN cells, which enabled us to identify molecularly distinct clusters of pMN NPCs, pre-motor neurons (pre-MNs) and pre-OPCs among other cell types (Scott et al., 2021) (Figure 1A). Among the genes that expressed transcripts highly enriched in pre-OPCs was *gsx2*, which encodes a homeobox transcription factor (Scott et al., 2021). The specific enrichment of *gsx2* transcripts in pre-OPCs at 36 hpf, combined with evidence from mice that Gsx2 regulates OPC formation (Chapman et al., 2018; Chapman et al., 2012), raised the possibility that Gsx2 functions in spinal cord pre-OPCs to guide OPC formation.

**Figure 1.**
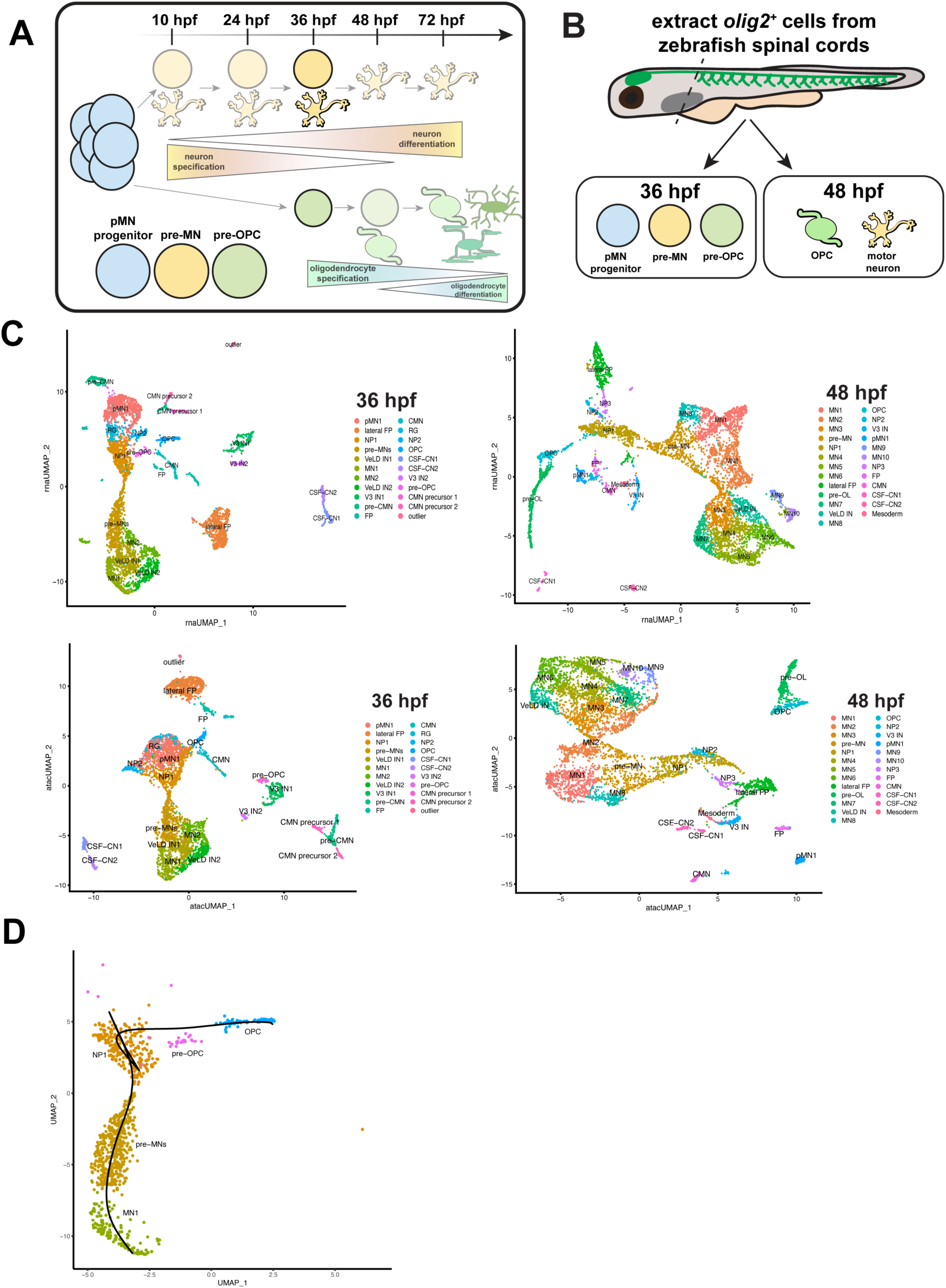
**scMulti-omics identify pre-OPCs as an intermediate cell state between neural progenitors and OPCs.** (A) Schematic of neuron and oligodendrocyte development in zebrafish. pMN progenitors (blue) produce neurons (yellow) from approximately 10-36 hpf and OPCs (green) beginning at 36 hpf, which mature as OLs by 72 hpf. (B) Schematic of cell isolation for scMultiome analysis. 36 and 48 hpf *Tg(olig2:egfp)* zebrafish larvae were decapitated and the trunk cells were dissociated and FAC sorted to isolate EGFP^+^ cells. Nuclei were extracted from these cells and submitted for sequencing. (C) UMAPs representing cell clusters generated by our scRNA-seq (top) and snATAC-seq (bottom) data at 36 (left) and 48 (right) hpf. (D) Slingshot trajectory analysis starting at NP1 and ending at either MN1 or OPC. Outlier cells were filtered for a maximum distance away from the center of each cluster. Black lines follow the predicted trajectories.

To investigate potential transcriptional mechanisms that regulate the transition from NPCs to OPCs, we generated a scMulti-omics dataset consisting of scRNA-seq and single-nuclei ATAC-sequencing (snATAC-seq) from zebrafish spinal cord cells at 36 and 48 hpf, the time at which OPCs are specified and begin to differentiate (Figure 1A, B). To capture NPCs, neurons, and OLCs specific to the pMN domain, we utilized *Tg(olig2:egfp)* zebrafish larvae, which express EGFP under control of *olig2* regulatory DNA (Shin et al., 2003) (Figure 1B). Similar to our prior scRNA-seq dataset (Scott et al., 2021), our clustering of expression data at 36 hpf using Seurat identified subpopulations of NPCs, pre-MNs, and pre-OPCs among other cell groups (Figure 1C). By 48 hpf, pre-OPCs and pre-MNs were absent from our cell clusters, presumably replaced by motor neurons, OPCs, and pre-myelinating oligodendrocytes (pre-OLs) (Figure 1C). We conducted trajectory analysis of the motor neuron and OPC lineages using Slingshot (Street et al., 2018) (Figure 1D) and visualized paths from NP1 to MN1 and OPC through their intermediate progenitors. These data identify pre-OPCs as a subset of NPCs that give rise to the OL lineage.

To further differentiate pre-OPCs from NP1s and pre-MNs, we used our scRNA-seq data to compare significant differential gene expression (log2FC >2 and adj p-value <0.05) across the three cell types at 36 hpf. All three cell types express both common genes and unique genes (Figure 2A). Examination of genes specific to NPCs, the motor neuron lineage, and the OL lineage revealed that NP1 cells most strongly express NPC genes, pre-MNs most strongly express motor neuron lineage genes, and pre-OPCs most strongly express a mix of NPC genes and OL genes (Figure 2B). To determine the differences in gene expression between pre-OPCs and NP1 and pre-MN clusters, respectively, we performed differential gene analysis (Figure 2C-E), which showed that pre-OPCs express *gsx2* at significantly higher levels than NP1 cells (Figure 2C). Further, *gsx2* transcripts were limited to pre-OPCs, with little expression evident in NP1 cells, the motor neuron lineage, or even OPCs (Figure 2E). Using our Slingshot analysis (Figure 1D), we plotted *gsx2* expression across pseudo time and observed that *gsx2* expression emerges during the transition from NP1 to pre-OPC and wanes as pre-OPCs transition to OPCs (Figure 2F). Together, these data support the possibility that Gsx2 functions within pre-OPCs as they transition from NP1 to OPC identities.

**Figure 2.**
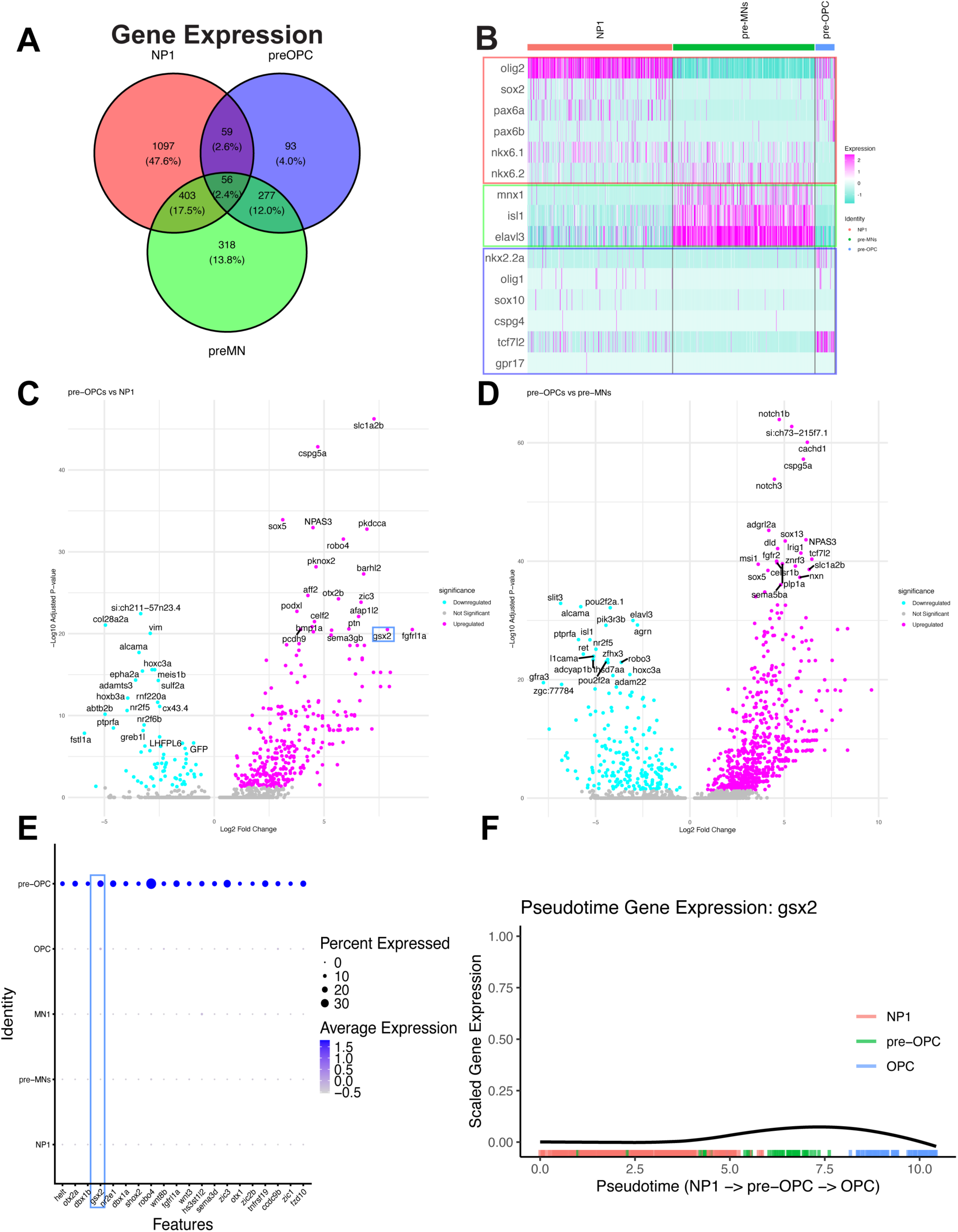
**pre-OPCs express *gsx2.*** (A) Venn diagram demonstrating the number of genes expressed in the clusters neural progenitor 1 (NP1), pre-OPCs, and pre-MNs from our 36 hpf scRNAseq dataset. (B) Heatmap of classically expressed neural progenitor genes (salmon, top), motor neuron genes (green, middle), and glial genes (blue, bottom) in NP1, pre-OPC, and pre-MN clusters. Magenta indicates high expression while turquoise indicates low expression. (C-D) Differential gene expression between pre-OPC and NP1 clusters (C) and pre-OPC and pre-MN clusters (D) at 36 hpf. Magenta dots indicate genes upregulated in pre-OPCs while turquoise dots indicate genes downregulated in pre-OPCs. Gray dots indicate non-significant genes. The most statistically significant upregulated and downregulated genes are labeled. (E) Dot plot representing the top differentially upregulated genes in pre-OPCs compared to all other clusters at 36 hpf. ∼20% of pre-OPCs express *gsx2* at 36 hpf. *gsx2* expression is outlined in blue. (F) Pseudotime analysis of *gsx2* expression from NP1 (salmon) to pre-OPC (green) to OPC (blue). *gsx2* expression rises as NP1 shifts to pre-OPC and fades as OPCs are specified. Overlap of clusters indicates transitions between cell states.

### Loss of *gsx2* results in precocious and excess OPCs

To investigate how *gsx2* might function in pre-OPCs, we used CRISPR/Cas9 genome editing to create loss-of-function alleles of *gsx2*. We recovered two alleles from independent founders and selected one, *co106*, for analysis. This allele has a 25 base-pair insertion and 2 base-pair deletion in exon 1, which are predicted to produce a premature stop codon and truncated protein (Figure 3A). *gsx2^−/−^* embryos and larvae were morphologically indistinguishable from wild-type siblings, consistent with scRNA-seq data across developmental time showing that *gsx2* expression is primarily limited to liver, primordial germ cells, and neural cells (Sur et al., 2023). To assess OPC formation, we fixed *gsx2^+/+^* and *gsx2^−/−^* larvae carrying the transgene *Tg(olig2:egfp)* at 36, 48, and 72 hpf, and 5 dpf. These timepoints encompass OPC specification (36 hpf), transition of OPCs to pre-OLs (48 hpf), initiation of OL differentiation and myelination (72 hpf) (Figure 1A), and mature OLs (5 dpf). We obtained transverse cryosections at the level of the trunk spinal cord and conducted immunohistochemistry to detect Sox10, which labels OLCs. At 36 hpf, Sox10^+^ OPCs were rarely evident in wild-type embryos (Figure 3B,C), consistent with our prior observations that *sox10* expression is first evident at about 40 hpf. By contrast, *gsx2* mutant embryos had a greater than two-fold excess of Sox10^+^ OPCs compared to their wild-type siblings (Figure 3B,C). Homozygous mutant embryos and larvae continued to have excess Sox10^+^ cells relative to wild-type siblings at each later stage of analysis (Figure 3B,C). Sox10^+^ OLC number plateaued at 48 hpf, suggesting that the excess number of OLCs results from excess production of OPCs from pMN progenitors, rather than from an elevated rate of OPC proliferation. Notably, at 3 dpf the excess number of OLCs was evident only in the ventral spinal cord (Figure 3D). These data suggest that in the absence of *gsx2* function, OPC specification begins prematurely and that excess pMN progenitors are selected for OLC fate.

**Figure 3.**
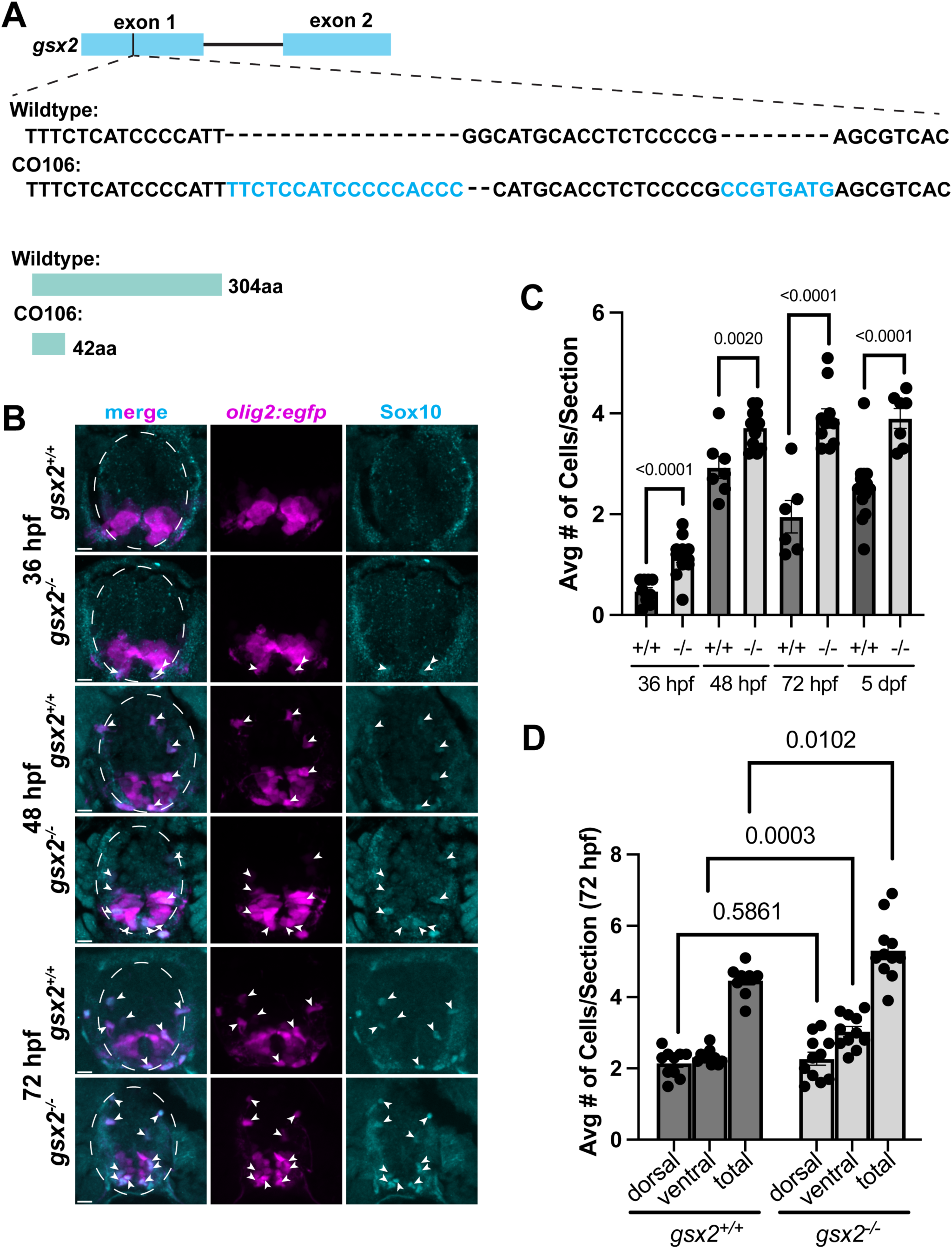
*gsx2* loss of function causes precocious and excess OPC formation. (A) Representation of CRISPR-Cas9 targeting to create *gsx2^co106^*, which consists of a 25 bp insertion and 2 bp deletion in exon 1 predicted to generate an early stop codon and truncated protein (teal). (B) Representative images of transverse sections of trunk spinal cord, dorsal up, obtained from *gsx2^+/+^* or *gsx2^−/^* embryos and larvae expressing EGFP (magenta) from the *olig2:egfp* transgene. Sections were processed to detect Sox10 expression (cyan). White dotted lines delineate the spinal cord. White arrows denote Sox10^+^ EGFP^+^ OLCs. (C) Graph showing the average number of OLCs per section at 36, 48, 72 hpf and 5 dpf. *n=* 10 larvae each for *gsx2^+/+^* and *gsx^−/−^* animals at each time point. (D) Graph showing the average number of OLCs per section at 3 dpf, with separation of dorsal and ventral cells. *n=* 10 larvae each for *gsx2^+/+^* and *gsx^−/−^* animals. For both C and D, each dot represents an individual zebrafish larva. P-values of unpaired *t-*tests are indicated on the plots. Error bars represent the mean with SEM. Scale bars: 5 µm.

To learn if loss of *gsx2* function affects spinal cord progenitors we performed immunohistochemistry to detect Sox2, a NPC marker, in combination with *olig2*:EGFP expression. At 36 and 48 hpf there were no obvious differences in Sox2 expression between *gsx2* mutant embryos and their wild-type siblings (Figure 4A). Additionally, there were no differences between the number of Sox2^+^ *olig2:*EGFP^+^ cells between genotypes at either timepoint (Figure 4B).

**Figure 4.**
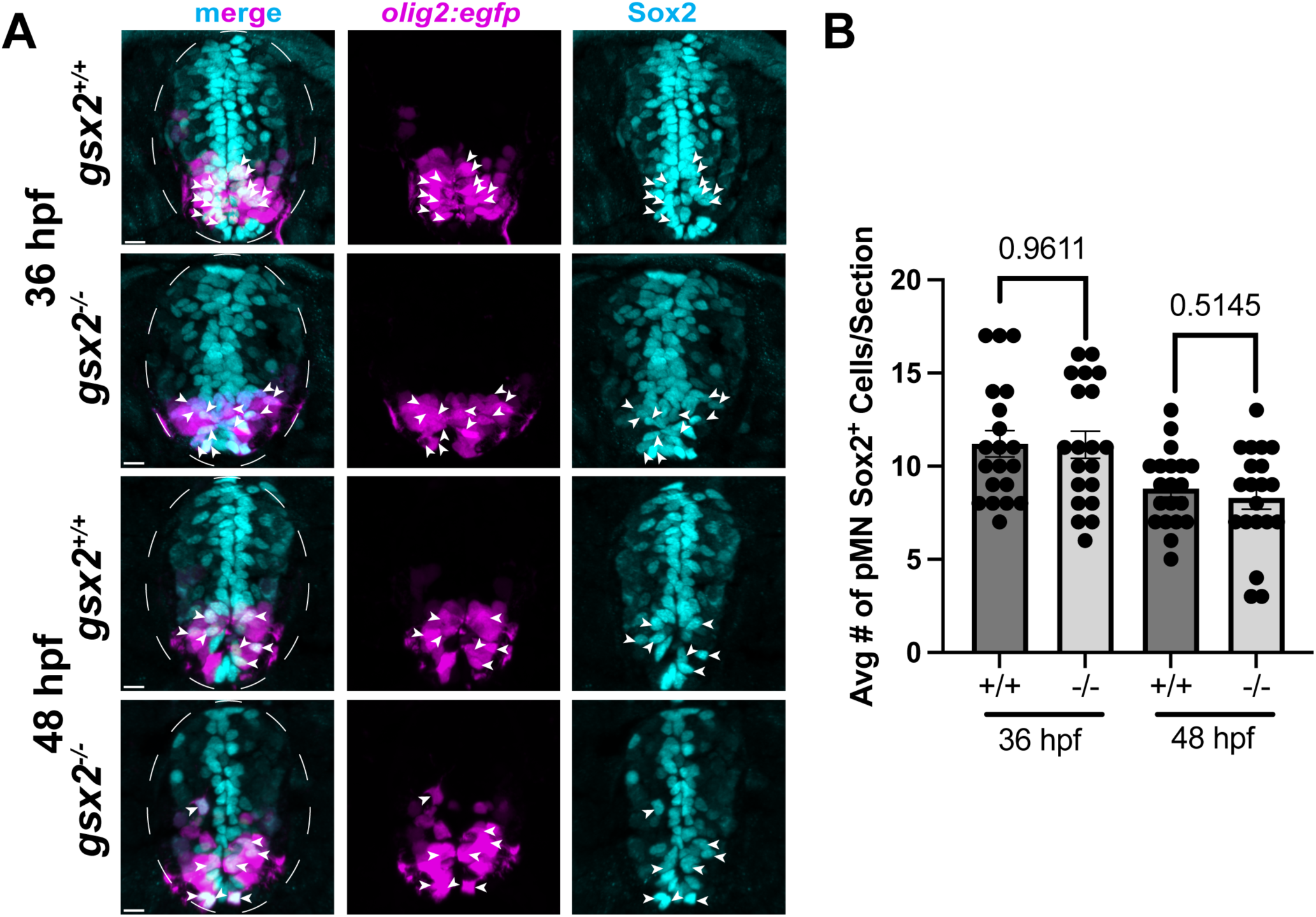
Loss of *gsx2* function does not alter neural progenitor cell number. (A) Representative images of Sox2 immunohistochemistry (blue) in transverse sections of *gsx2^+/+^*or *gsx^−/−^* larvae expressing *olig2:egfp* (magenta) at 36 and 48 hpf. White dotted lines delineate the spinal cord. White arrows denote Sox2^+^/*olig2*^+^ cells. (B) Graph showing average number of Sox2^+^/*olig2*^+^ cells per section at 36 and 48 hpf. *n=* 20 larvae for *gsx2^+/+^* or *gsx^−/−^* larvae at each timepoint. Each dot represents an individual zebrafish larva. P-values of unpaired *t-*tests are indicated on the plots. Error bars represent the mean with SEM. Scale bars: 5 µm.

### Loss of *gsx2* function increases the number of myelinating oligodendrocytes but does not result in premature differentiation

pre-OPCs but not OPCs express *gsx2* (Figure 2E), therefore, we hypothesized that *gsx2* function is limited to formation but not differentiation of the oligodendrocyte lineage. To test this, we assessed OLC differentiation by using in situ RNA hybridization to detect transcripts encoded by *myrf*, a marker of pre-myelinating OLs, and *mbpa*, which marks myelinating OLs. At 48 hpf, despite the excess number of Sox10^+^ OLCs in mutant embryos (Figure 3C), there was no difference in the number of *myrf^+^* or *mbpa^+^* cells in *gsx2^-/-^* embryos compared to wild-type siblings (Figure 5A-C). By contrast, at 72 hpf, *gsx2^-/-^* larvae had more *myrf^+^* and *mbpa^+^* cells than wild-type siblings (Figure 5D-F). To assess the ratio of OLs to all OLCs, we simultaneously detected *sox10* and *mbpa* transcripts. Consistent with our immunohistochemistry experiments, *gsx2^-/-^*larvae had more *sox10^+^* OLCs than wildtype. However, there was no significant difference in the ratio of *sox10^+^/mbpa^+^*cells between genotypes (Figure 5I). These data indicate that loss of *gsx2* function changes neither the timing of OL differentiation nor the fraction of OLCs that differentiate as OLs. We interpret these data to mean that Gsx2 regulates OPC formation but not subsequent differentiation.

**Figure 5.**
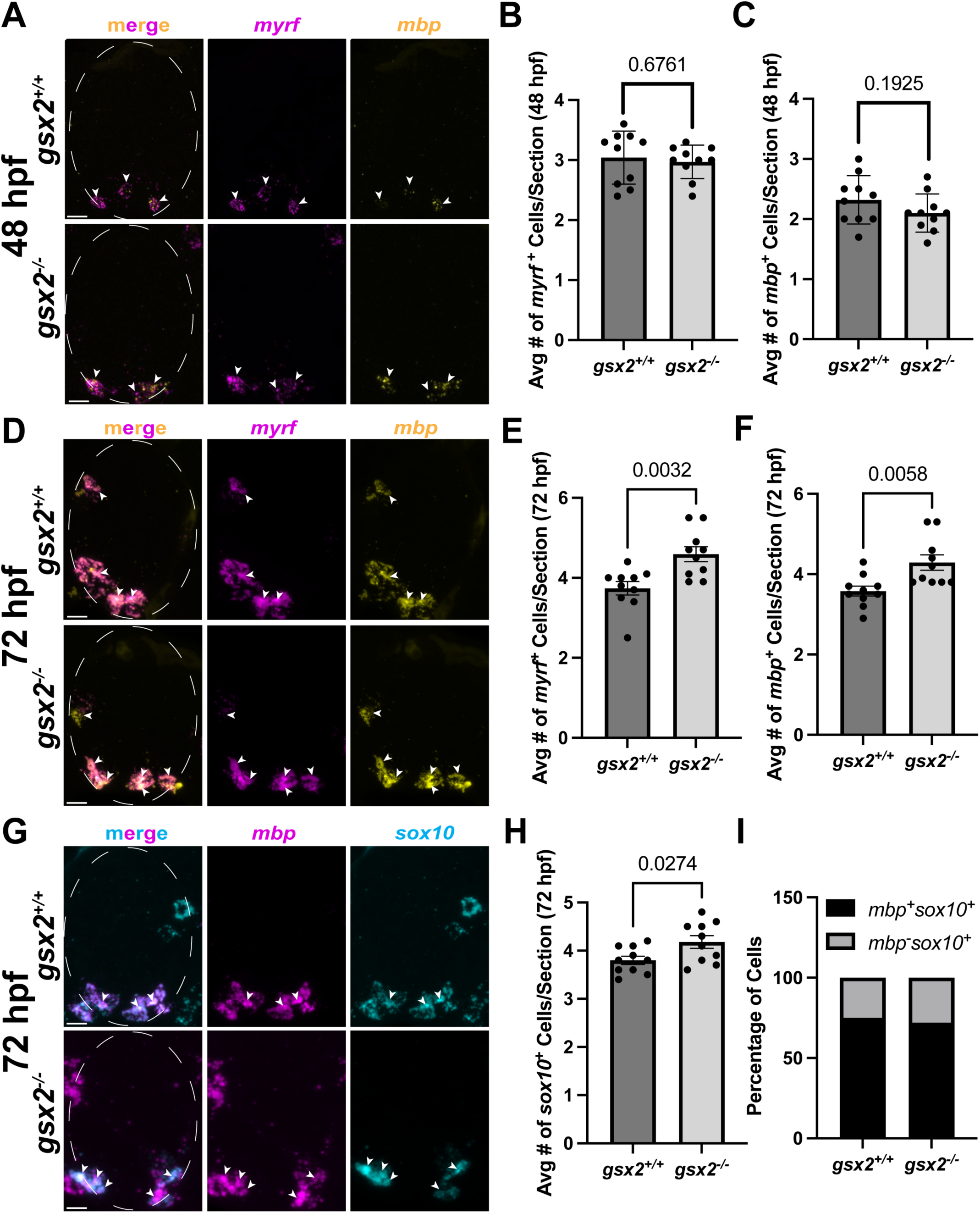
**Loss of *gsx2* function does not affect oligodendrocyte differentiation.** (A,D,G) Representative images of transverse sections through the trunk spinal cord processed by in situ RNA hybridization to detect *myrf* (magenta) and *mbpa* transcripts (A,D) or *mbpa* (magenta) and *sox10* (cyan) transcripts (G) in *gsx2^+/+^* and *gsx^−/−^* larvae. White dotted lines delineate the spinal cord. White arrows denote cells that express both transcripts. (B,C) Graphs showing the average number of *myrf*^+^ (B) and *mbpa*^+^ (C) cells per section at 48 hpf. (E,F) Graphs showing the average number of *myrf*^+^ (E) and *mbpa*^+^ (F) cells per section at 72 hpf. For B, C, E, F, and H, each dot represents an individual zebrafish larva, *n= 10* larvae each for *gsx2^+/+^* and *gsx2^-/-^* larvae. P-values of unpaired *t-*tests are indicated on the plots. Error bars represent the mean with SEM. (I) Percentage of cells that were *sox10^+^/mbpa^+^* vs *sox10^+^/mbp^−^* in *gsx2^+/+^* and *gsx^−/−^* larvae. Scale bars: 5 µm.

### Single-cell muti-ome analysis reveals potential candidate regulators of *gsx2* expression

To uncover factors that might activate *gsx2* transcription we identified genes differentially expressed in pre-OPCs compared to NP1 cells. We then used a Pearson correlation to determine if the expression of each gene was positively correlated with *gsx2* expression (Figure 6A). Our analyses identified transcription factor encoding genes *zic1, zic3, and helt, hs3st1l2,* which encodes a heparan sulfate sulfotransferase that modulates FGF, Shh, and Wnt signaling, and Wnt regulators *fzd10* and *lef1*. To identify potential repressors of *gsx2* that function at the pre-OPC to OPC transition we performed differential expression analysis to reveal genes that are highly expressed in OPCs compared to pre-OPCs. We then similarly used a Pearson correlation to determine which of these genes are negatively correlated with *gsx2* expression (Figure 6B). Among the most highly expressed in OPCs and negatively correlated with *gsx2* were transcription factor-encoding genes *nkx2.2a, sox6, sox9a,* and *myrf* (Figure 6B).

**Figure 6.**
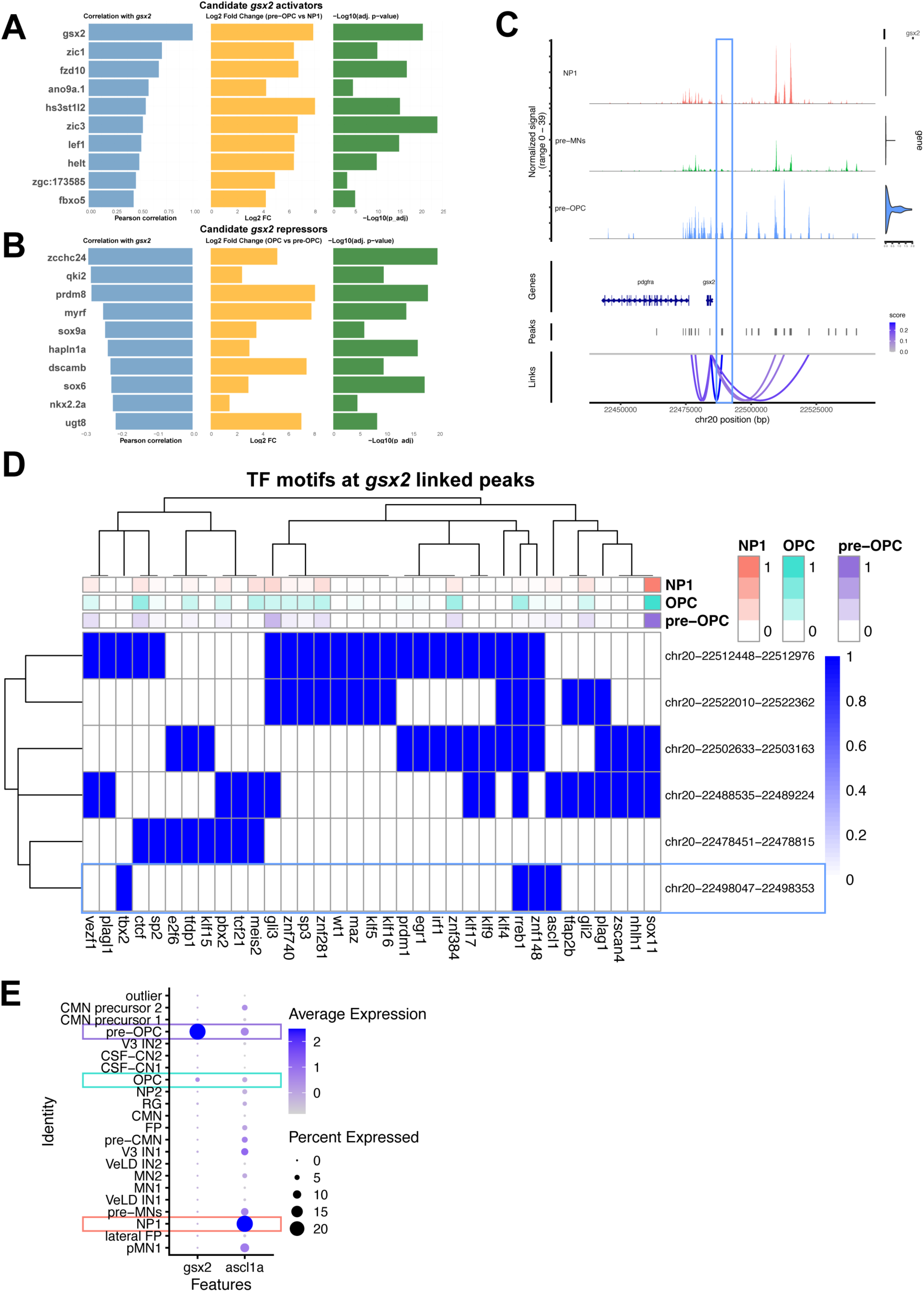
Identification of potential regulators of *gsx2* expression. (A) Genes expressed at high level in pre-OPCs relative to NP1 cells and positively correlated with *gsx2* expression. (B) Genes expressed at low level in OPCs relative to pre-OPCs and negatively correlated with *gsx2*. For (A) and (B), Pearson correlation is shown in blue, differential expression is shown in yellow, and p-value is shown in green. (C) Linked peak analysis of *gsx2* at 36 hpf for NP1 (pink), pre-MNs (green), and pre-OPCs (blue). Violin plots of gsx2 expression (right). Peaks (middle) indicate regions of chromatin accessibility and map to genomic locations (bottom). Individual peaks linked with gene expression are indicated by blue lines, with darker blue representing a stronger correlation with expression. The blue box outlines a peak strongly linked to gene expression. (D) Heatmap of motifs corresponding to transcription factor binding sites at the six peaks linked with gene expression from (C) at the *gsx2* locus. Expression of each transcription factor-encoding gene in NP1, pre-OPC, and OPC clusters is indicated above the heatmap. The blue box highlights the peak having accessibility most highly correlated with gsx2 expression, as indicated by the dark blue line in (C). (E) Dotplot of *gsx2* and *ascl1a* expression across all clusters at 36 hpf, highlighting pre-OPC (purple), OPC (cyan), and NP1 clusters (orange).

To more rigorously identify potential regulators of *gsx2* expression, we explored how gene expression correlates to chromatin accessibility at the *gsx2* locus. We identified chromatin peaks, which represent regions of accessible chromatin, within 20 kb spanning the *gsx2* locus. We then performed a linked-peak analysis, which predicts which of these chromatin peaks might influence *gsx2* expression by correlating peak accessibility with gene expression (Figure 6C). This analysis identified six peaks with highly correlated accessibility and *gsx2* gene expression in pre-OPCs, suggesting that these regions might harbor cis-regulatory elements that control *gsx2* transcription. Similar peaks were not evident in NP1 and pre-MN clusters (Figure 6C). To investigate potential regulators of *gsx2,* we identified transcription factor motifs present at the six peaks linked to *gsx2* expression. We then examined the expression of genes encoding the transcription factors associated with each motif in pre-OPCs (Figure 6D). Genes encoding transcription factors uncovered by this analysis include *sox11, gli2, gli3, znf740, znf384, znf281, ctcf,* and *ascl1a.* NP1 cells, the apparent progenitors of pre-OPCs, express *ascl1a* at a high level (Figure 6E), highlighting Ascl1 as a potential activator of *gsx2* expression, consistent with evidence that Ascl1 promotes oligodendrocyte development (Parras et al., 2007; Sugimori et al., 2007; Tran et al., 2023; Vue et al., 2014; Winkler et al., 2020).

### A potential Gsx2-centered gene regulatory network for spinal cord OPC formation

Our mutant analyses showed that loss of *gsx2* alters the timing of OPC formation (Figure 3). To uncover the mechanisms through which *gsx2* might regulate OPC timing, we utilized Pando (Fleck et al., 2023) to infer a gene regulatory network (GRN) of transcription factors and target genes downstream of Gsx2 from integration of our gene expression and chromatin accessibility data. Our predicted GRN identified several genes known to be involved in OPC specification and OLC differentiation immediately downstream of *gsx2* expression (Figure 7A). We next identified the preferred Gsx2 DNA-binding sequence using the JASPAR database (Figure 7B). Using our snATAC-seq dataset, we then identified all the open chromatin regions in all our 36 hpf clusters that contain the Gsx2 motif, revealing 9,422 peaks. We filtered those peaks for those that are accessible in pre-OPCs, narrowing our chromatin regions to 4,882 peaks. Linking the snATAC-seq peaks with the scRNA-seq gene expression, we determined which of these pre-OPC-accessible peaks were linked to genes that are expressed in pre-OPCs, finding 1,484 peaks. Finally, we filtered these peaks to find those that were linked to genes that were differentially expressed in pre-OPCs compared to all other clusters. Our results revealed 83 peaks that contain the Gsx2 DNA-binding motif, are accessible in pre-OPCs, and are linked to genes that are differentially expressed in pre-OPCs (Table 1). These 83 peaks represent predicted targets of Gsx2 binding (Figure 7C), several of which are known to be important for OLC development and are downstream of *gsx2* in our GRN (Figure 7A). Common factors between the two predictions were *sox5* and *tcf7l2,* which encode transcription factors that regulate oligodendrocyte linage cell development (Hammond et al., 2015; Stolt et al., 2006; Weng et al., 2017; Zhang et al., 2021; Zhao et al., 2016). We further explored the expression of the predicted targets with the lowest p-values across all our clusters and found that all have high expression in pre-OPCs, almost no expression in our neuronal clusters, and moderate expression in our neural progenitor clusters (Figure 7D). Together, our *in vivo* and multi-omics results suggest that *gsx2* is briefly expressed in pre-OPCs to coordinately time expression of target genes that promote OPC specification.

**Figure 7.**
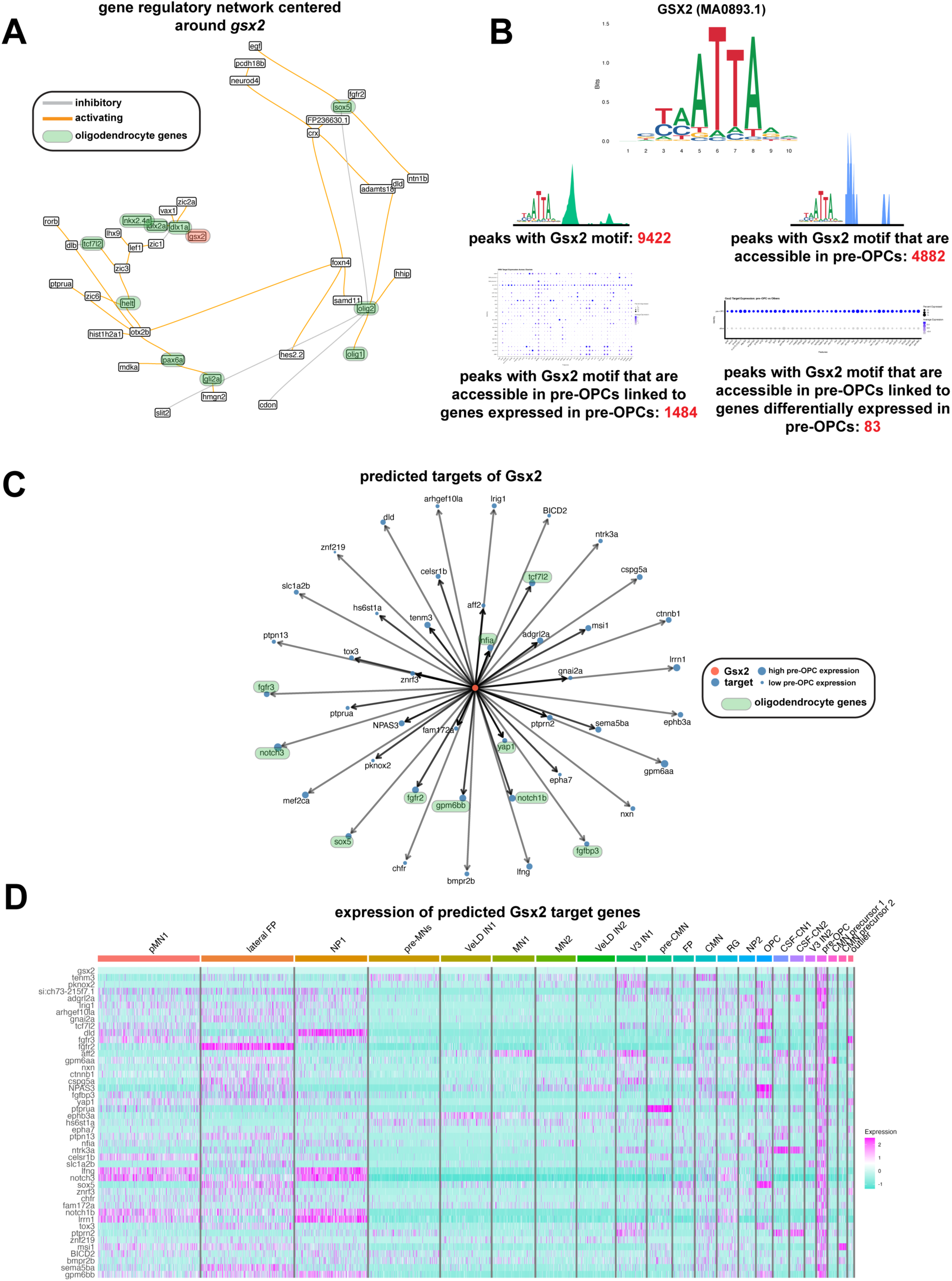
**Exploration of a *gsx2*-centered gene regulatory network**. (A) Linked peak analysis of *gsx2* at 36 hpf for NP1 (pink), pre-MNs (green), and pre-OPCs (blue). Gene expression of *gsx2* (right) is highest in pre-OPCs. Peaks (middle) indicate regions of chromatin accessibility and map to genomic locations (bottom). Individual peaks linked with gene expression are indicated by blue lines, with darker blue representing a stronger correlation with expression. The blue box outlines peaks strongly linked to gene expression. (B) A predicted gene regulatory network centered around *gsx2* at 36 hpf. Transcription factors highlighted in green are known to be involved in OPC specification. Orange lines indicate activating relationships whereas gray lines indicate inhibitory relationships. (C) Logo for the Gsx2 motif (JASPAR) (top). Visualizations of analyses conducted using our scRNAseq and snATACseq 36 hpf datasets. For our entire 36 hpf dataset, we found 9422 peaks (regions of open chromatin) with the Gsx2 motif (preferred binding sequence) (top left). Of those peaks, we found that 4882 of them are peaks that are accessible in pre-OPCs (top right). We analyzed those peaks accessible in pre-OPCs and found 1484 were linked to genes that are expressed in pre-OPCs (bottom left). Of those genes, we found that 83 are differentially upregulated in pre-OPCs (bottom right). (C) A map highlighting predicted targets of Gsx2 (top statistically significant genes that are differentially upregulated in pre-OPCs, have peaks that are accessible in pre-OPCs, and whose peaks possess the Gsx2 motif). Genes highlighted in green are known to be involved in the oligodendrocyte lineage. Blue dots represent target genes while the center orange dot represents Gsx2. Larger dots indicate higher expression in pre-OPCs whereas smaller dots indicate lower expression. (D) Heatmap of the predicted Gsx2 target genes across all clusters a 36 hpf. Magenta indicates high expression while turquoise indicates low expression.

## DISCUSSION

Whereas many investigations have uncovered mechanisms that regulate differentiation of oligodendrocyte lineage cells (Emery and Wood, 2024) a major gap has persisted in our understanding of mechanisms that specify neural progenitors for oligodendrocyte lineage cell fate. This gap exists, in large part, because of our incomplete knowledge of the specific neural progenitors that give rise to OPCs and the gene regulatory mechanisms that guide their transition. With this work and a prior study (Scott et al., 2021) we have identified pre-OPCs in the developing zebrafish spinal cord as a transition state between neural progenitors and OPCs. pre-OPCs, also called pri-OPCs, similarly have been described in mice and humans (Fu et al., 2021; Huang et al., 2020; Marques et al., 2018; Van Bruggen et al., 2022; Weng et al., 2019; Winkler et al., 2020). Together these studies provide a path toward elucidating transcriptional control mechanisms that initiate oligodendrocyte development.

To begin interrogating regulatory mechanisms of OPC specification, we chose to test the function of the homeobox transcription factor encoding gene *gsx2*. Prior work using mice had already shown that loss of *Gsx2* function alters oligodendrocyte development, giving us an opportunity to assess if similar mechanisms operate across different species in OPC specification. Our trajectory analysis suggested a cell lineage progressing from neural progenitors (NP1) through pre-OPCs to OPCs. Mapping *gsx2* expression on to this lineage indicated that pre-OPCs but not NP1 progenitors or OPCs express *gsx2*. Additionally, *gsx2* transcripts were only evident in about 20% of pre-OPCs. One interpretation of these data is that many pre-OPCs express *gsx2* at low levels insufficient for detection by our scRNA-seq methods. Alternatively, pre-OPCs might express *gsx2* only briefly as they progress through pre-OPC state. In either case, these data are consistent with the possibility that a small burst of *gsx2* expression contributes to a mechanism that controls a NP1 to pre-OPC to OPC transition.

Our gene function analyses revealed that *gsx2* mutant larvae produced OPCs prematurely and in excess number, strikingly similar to OPC formation in the forebrain of *Gsx2* mutant mice (Chapman et al., 2018; Chapman et al., 2012). Despite their early formation, oligodendrocyte differentiation occurred on schedule in *gsx2* mutant zebrafish larvae. In mice, the elevated numbers of OPCs formed in the absence of *Gsx2* function was accompanied by an apparent deficit in neurogenesis, suggesting that *Gsx2* helps switch neural progenitors from a neurogenic program to gliogenesis (Chapman et al., 2012). In zebrafish, the number of neural progenitors appeared to be unaffected by loss of *gsx2* function, but the number of pre-OPCs relative to neural progenitors and neurons is so small that it is difficult for us to identify any change in neural cell fate coupled with excess OPC formation. Nevertheless, our data support a model in which Gsx2 momentarily holds cells in a pre-OPC state. Mechanisms that downregulate *gsx2* transcription are therefore likely important for OPC specification.

The highly specific temporal and cell type expression of *gsx2* led us to evaluate mechanisms that might regulate its transcription. Our snATAC-seq identified regions of open chromatin near *gsx2* potentially occupied by transcription factors and chromatin regulators that pre-OPCs express. For example, we identified potential binding sites for Ascl1, which, in mice, promotes OPC formation (Parras et al., 2007; Sugimori et al., 2007; Tran et al., 2023; Vue et al., 2014). Ascl1 also promotes neurogenesis, but driving Gsx2 expression can suppress the neurogenic function of Ascl1 through physical interaction (Roychoudhury et al., 2020). This raises the possibility that Ascl1-driven expression of Gsx2 in pre-OPCs helps switch Ascl1 function from neurogenic to gliogenic. We also found potential binding sites for Gli2 and Gli3, which mediate transcription activity in response to Shh signaling. Subsequent to patterning the spinal cord dorsoventral axis by Shh (Dessaud et al., 2008), a transient elevation of Shh signaling activity promotes OPC formation (Danesin et al., 2006; Danesin and Soula, 2017; Jiang et al., 2017; Oustah et al., 2014; Scott et al., 2020). Notably, in the forebrains of mice Shh drives formation of *Gsx2*^+^ tripotential progenitors that give rise to oligodendrocytes in addition to astrocytes and olfactory bulb interneurons (Zhang et al., 2020) and CUT&Tag-seq experiments revealed Gli3 binding near *Gsx2* (Gao et al., 2025). Therefore, *gsx2* could be a direct target of Shh signaling in OPC specification. The striking loss of *gsx2* transcripts that marks the pre-OPC to OPC transition suggests that transcriptional inhibition of *gsx2* also contributes to OPC specification. Consistent with this possibility, we identified potential binding sites for Nkx2.2. In mice, Nkx2.2, which promotes oligodendrocyte development, represses *Pdgfra* transcription by binding regulatory DNA at the *Pdgfra* locus (Zhu et al., 2014).

Our analysis additionally revealed numerous genes that function in oligodendrocyte development that might be regulated by Gsx2. Among these are *fgfr2* and *fgfr3*, *notch1b* and *notch3, sox5, nfia,* and *tcf7l2*. Pharmacological inhibition of Fgfr signaling in early neural development blocked OPC specification (Farreny et al., 2018) and *notch3* mutant zebrafish larvae had a transient deficit of OPCs (Zaucker et al., 2013). In mice, loss of *Sox5* function caused precocious spinal cord OPC formation and elevated OPC numbers (Stolt et al., 2006), similarly to loss of *gsx2* function in zebrafish. Notably, excess OPCs accumulated in the ventral spinal cord of both *Sox5* mutant mice (Stolt et al., 2006) and *gsx2* mutant zebrafish, raising the possibility that these factors also regulate OPC migration to dorsal spinal cord. Nfia promotes the transition from neurogenesis to gliogenesis (Deneen et al., 2006) and Tcf7l2 promotes oligodendrocyte differentiation (Hammond et al., 2015; Weng et al., 2017; Zhang et al., 2021; Zhao et al., 2016). Altogether, these observations are consistent with the possibility that Gsx2 coordinates expression of genes that regulate the timing of OPC specification and initiate oligodendrocyte differentiation.

The similarity of *Gsx2/gsx2* mutant phenotypes in mouse forebrain and zebrafish spinal cord indicates that Gsx2 functions across species and nervous system regions to regulate OPC specification. In support of this possibility, an analysis of scRNA-seq data from human fetal forebrain identified pre-OPCs that express *GSX2* and placed GSX2 function within a regulatory network guiding the pre-OPC to OPC transition (Van Bruggen et al., 2022). We therefore propose that the highly regulated, transient expression of *GSX2* orthologs contributes to a gene regulatory network that determines the timing of the pre-OPC to OPC transition.

## METHODS

### Zebrafish lines and husbandry

All animal work was approved by the Institutional Animal Care and Use Committee (IAUCUC) at the University of Colorado School of Medicine. All non-transgenic embryos were obtained from pairwise crosses of males and females from the AB strain. All transgenic embryos were obtained from pairwise crosses of males or females from the AB strain to males or females from the *Tg(olig2:egfp)^vu12^* strain (Shin et al., 2003). Embryos were raised in petri dishes at 28.5°C in egg water (6g Instant Ocean in 20L miliQ water) and staged according to hours post fertilization (hpf) and morphological features (Kimmel et al., 1995). Sex cannot be determined at zebrafish embryonic and larval stages. For embryo injections, eggs were collected within 30 minutes of breeding and injected at the single-cell stage. Larvae fixed for experiments were euthanized with 4% Tricaine mesylate prior to whole fixation in 4% paraformaldehyde in 1X PBS (PFA). When larvae required genotyping, tail clips were taken from fixed larvae and lysed.

### CRISPR-Cas9 mutagenesis

We utilized the CRISPR-Cas9 system to generate our *gsx2* loss-of-function mutants. We used Integrated DNA Technologies (IDT) to select the top two predesigned Alt-R CRISPR-Cas9 guide RNAs (gRNAs) that target the coding sequence of exon 1 for *gsx2* (see key resources table for gRNA sequences). To prepare our gRNAs and synthesize the ribonucleoprotein (RNP) complex, we followed the “Alt-R CRISPR-Cas9 System: *in vitro* cleavage of target DNA with ribonucleoprotein complex” protocol (https://sfvideo.blob.core.windows.net/sitefinity/docs/default-source/protocol/alt-r-crispr-cas9-protocol-in-vitro-cleavage-of-target-dna-with-rnp-complex.pdf?sfvrsn=88c43107_2, 2017). The RNP complex was created immediately prior to injections by combining 0.5 µg Cas9 (IDT) diluted in Cas9 working buffer (20 mM HEPES, 150 mM KCl, pH 7.5) with 100 ng of each *gsx2* gRNAs and incubating the complex at 37°C for 10 minutes. The RNP complex was kept at room temperature until injected, where 2-3 nL was injected into wild-type embryos at the single-cell stage. F0 injected larvae were screened for insertions and/or deletions created by DNA cutting and repair with PCR primers designed specifically for the gRNA target site (see key resources table for PCR primers and product sequences). F0 larvae were raised to adulthood and individually outcrossed to wild-type fish. F1 larvae were anesthetized and genomic DNA was extracted using 50 mM NaOH followed by 1M Tris, pH 9.0 neutralization. Several individual F1 animals from at least two unique F0 founders were screened for mutant alleles using the same PCR primers described above. Mutant alleles were sequenced and aligned with the published genome to identify function-breaking mutation. F1 adults carrying the selected *gsx2^CO106^* mutant allele were outcrossed to wild-type fish and the F2 larvae were raised for experiments. All wild-type and mutant larvae were genotyped and alleles were confirmed prior to their use in experiments.

### Immunohistochemistry

Larvae were fixed at indicated time points in 4% PFA, rocking at room temperature for 3 hours. Larvae were rinsed in 0.1% Triton/1x PBS (PBSTx) and stored at 4°C. Larvae were embedded in 1.5% agar/30% sucrose overnight at 4°C. Blocks were frozen on dry ice and 15 µm transverse sections were taken with a cryostat microtome and collected on polarized slides. Slides were mounted in Sequenza racks (Ted Pella cat 36105), washed 3 x 5 minutes with PBSTx, and blocked for 1 hour in 2% goat serum/2% bovine serum albumin/PBSTx before being placed in primary antibody (mixed in block) overnight at 4°C. Primary antibodies used in this paper include: rabbit anti-Sox10 (1:500, (Park et al., 2005)) and rabbit anti-Sox2 (1:500, Abcam). Sections were washed for 1.5 hours in PBSTx, then incubated in secondary antibody (1:250, in block) for 2 hours at room temperature. Secondary antibodies used in this paper include: AlexaFluor 568 goat anti-rabbit (cLife Technologies). Sections were washed for 1 hour continuously in PBSTx and then mounted in Vectashield (Vector Laboratories, H-1000-10). For imaging analysis, fixed zebrafish sections were acquired using a 20x plus optovar 1.6x objective on a Zeiss Cell Observer SD 25 spinning disk confocal system (Carl Zeiss). Z stacks images were captured on Zen software (Carl Zeiss) and were processed and analyzed using

Fiji/ImageJ. For imaging analysis with the Sox2 antibody, fixed zebrafish sections were acquired using an Andor Dragonfly 200 40µm Spinning Disk Confocal (Oxford Instruments) and DMi8 inverted microscope (Leica) with 40x/na1.3 HC PL APO oil objective. Z stack images were captured on Fusion v2.4.0.14 software (Oxford Instruments) and were processed and analyzed using Fiji/ImageJ.

### Fluorescent *in situ* RNA hybridization

Fluorescent in situ RNA hybridization was performed using RNAscope Multiplex Fluorescent V2 Detection Kit (ACD 323110). Zebrafish embryos were fixed at either 2 dpf or 3 dpf with 4% PFA, embedded as described above, and 12 µm cryosections were processed using manufacturers protocol and synthesized probes for *sox10* Ch2, *mbpa* Ch3 or *myrf* Ch2 followed by Opal dyes 570 (Akoya FP1488001) or 650 (Akoya FP1496001). For imaging analysis, Z stack images were acquired from fixed zebrafish sections using Zen software (Carl Zeiss) with 20x objective on Zeiss LSM 880. Images were analyzed in Imaris Viewer software (version 10.2). 10 sequential images were captured for each larva at each genotype and averaged per larvae for statistical analyses.

### Quantifications/Statistical Analyses

#### Image Blinding

The genotypes of fixed slides were blinded prior to imaging and were unblinded once analysis was complete.

#### Cell Counts

Cells were counted in transverse sections of *Tg(olig2:egfp)* larvae processed for immunohistochemistry as described above to detect Sox10 or Sox2. For Sox10 experiments, the number of dorsal and ventral Sox10^+^ cells from 10 sequential sections were counted per larva. Only Sox10^+^ cells that were also *olig2^+^* were counted. The number of Sox10^+^ cells per larva were determined by averaging dorsal, ventral, and total numbers across the 10 sequential sections. A minimum of 10 larvae with 10 sequential sections each were used for each experimental group. For Sox2 experiments, the number of Sox2^+^ cells were counted from 10 sequential sections per larva and averaged for total numbers per larva as described above. Only Sox2^+^ cells that were in the *olig2^+^*pMN domain were counted.

#### Statistical Analysis

All data analysis, statistics, and plotting were performed in GraphPad Prism (version 10). *p* values involving only two groups were calculated using an unpaired two-tailed *t* test. Significance levels were determined using a confidence interval of 95%. The data in the plots and in the text are presented as means +/-SEM. A minimum of 10 individual larvae were used in all quantified studies.

#### Cell dissociation and nuclei isolation

36 and 48 hpf embryos were dechorionated and screened for *Tg(olig2:egfp)* expression using a fluorescent stereomicroscope. 400 embryos for each timepoint were euthanized with 4% Tricaine MS-222 and decapitated using a razor blade. The embryo trunks were transferred to 1.7 mL microcentrifuge tubes in groups of 50 on ice. To remove the yolk, we added 100 µL of pre-cooled calcium-free Ringer’s solution (1.16 mL 5M NaCl, 130 µL 1M KCl, 250 µL 1M HEPES pH 7.3, 48.445 mL PBS). Trunks were pipetted up and down gently with a p200 pipette for 15 minutes on ice. After an additional 5 minutes 500 µL of 10 mg/mL protease solution was added (312.5 µL DNase, 125 µL EDTA, 24.563 mL 1x PBS (-CaCl, -MgCl), 250mg protease from Bacillus licheniformis). The tissue was incubated on ice for 15 minutes while gently homogenizing every 3 to 5 minutes with a p200. Next, 200 µL of stop solution (15 mL FBS, 200 µL 1M CaCl2, 5 mL 10x PBS) was added, mixed, and spun down for 5 minutes at 400g at 4°C. Supernatant was removed, leaving around 100 µL to not disturb the pellet. 1 mL of chilled suspension solution (500 µL 1M CaCl_2_, 500 µL 100x pen strep, 500 µL FBS, 45 mL Leibovitz Medium Solution) was added, and the pellet was fully resuspended and spun down at 400g for 5 minutes at 4°C. Supernatant was removed, leaving around 100 µL to not disturb the pellet. 400 µL of chilled suspension solution was added, the pellet was fully resuspended, and the solution was filtered through a 35 µM strainer into a 1.7 mL microcentrifuge tube. Four sample preps were pooled into one tube, resulting in two filtered tubes per time point. The samples were FAC sorted to distinguish EGFP^+^ cells using a MoFo XDP100 cell sorter at the CU-SOM Cancer Center Flow Cytometry Shared Resource. Sorted samples were collected in 1.7 mL FBS coated microcentrifuge tubes.

To isolate nuclei from our FAC-sorted EGFP^+^ cells, we followed the nuclei isolation protocol from 10x genomics (https://cdn.10xgenomics.com/image/upload/v1660261285/support-documents/CG000124_Demonstrated_Protocol_Nuclei_isolation_RevF.pdf, 2023). We spun the cells down at 900G for 5 minutes at 4°C. The resulting supernatant was discarded, being careful not to disturb the pellet. We added 100 µL of 0.1x lysis buffer and gently resuspended the pellet by pipetting. The samples were incubated on ice for 1 minute after which 1 mL of ice-cold wash buffer was added to lysed cells and mixed by pipetting. Samples were spun down at 900g for 10 minutes at 4°C. The supernatant was discarded, being careful not to disturb the pellet. The wash buffer and centrifuge steps were repeated twice. After the last spin, cells were resuspended in 30 µL of diluted nuclei buffer. The nuclei were then immediately transferred on ice to the CU-SOM Genomics Shared Resource where they were counted and submitted for quality control and subsequent sequencing.

#### Multi-omics sequencing (10x)

Fastq sequencing files from 10 X Genomics multiomic single-nuclei RNA-seq and ATAC-seq were processed through Cell Ranger ARC (v1.0.1) with a zebrafish GRCz11 library to obtain UMI gene expression counts and ATAC peak fragment counts. These were analyzed using the standard methods in the Signac (v1.12.0) package (Stuart et al., 2021) in R. Briefly, a Seurat Object was created from the matrix.h5 and fragments.tsv.gz files, annotated with GRCz11.v99.EnsDb, and ATAC peaks corrected by calling with macs2 (v2.2.7.1) and finally filtered to exclude cells with gene count <750, total ATAC counts <300, percent mitochondrial gene expression >10%, nucleosome expression >2 or TSS enrichment <1. Gene expression was normalized with SCTransform and the dimensionality was reduced with PCA. DNA accessibility was processed by performing latent semantic indexing. The Seurat weighted nearest neighbor method was used to compute a neighbor graph which was visualized with UMAP and clusters were annotated based on the expression of marker genes. A pseudotime trajectory through the NP1, pre-OPC, OL and pre-MN clusters was calculated with Slingshot (Street et al., 2018). The *gsx2* centered GRN was calculated with Pando (Fleck et al., 2023).

#### Bioinformatics analyses

https://github.com/kimarena16/gsx2_2025.git

#### Author Contributions

K.A. and B.A. designed research, K.A., C.K, M.A., and B.A. performed research, K.A., R.O., and C.K. analyzed data, and K.A. and B.A. wrote the paper, K.A., B.A., S.F., and C.S. funded the studies.

## Acknowledgments

This work was supported by the National Institutes of Health (NIH)/National Institute of Neurological Disorders and Stroke (R01 NS124166 to S.J.F., B.H.A., and C.G.S.; R35 NS122191 to B.H.A; and F32 NS134612 to K.A.A.), and a gift from the Gates Frontiers Foundation to B.H.A. We thank Dr. Tyler Gibson for technical advice for bioinformatics analyses and zebrafish facility staff for animal care. We also thank Christine Childs, MT (ASCP) and Dmitry Baturin, MD PhD at the SOM Cancer Center Flow Cytometry Shared Resource and support grant from NCI P30CA046934. Finally, we thank Okyong Cho, MS at the University of Colorado Anschutz Medical Campus Cancer Center Genomic Shared Resource Core Facility.

The authors declare no conflict of interest.

## KEY RESOURCES TABLE

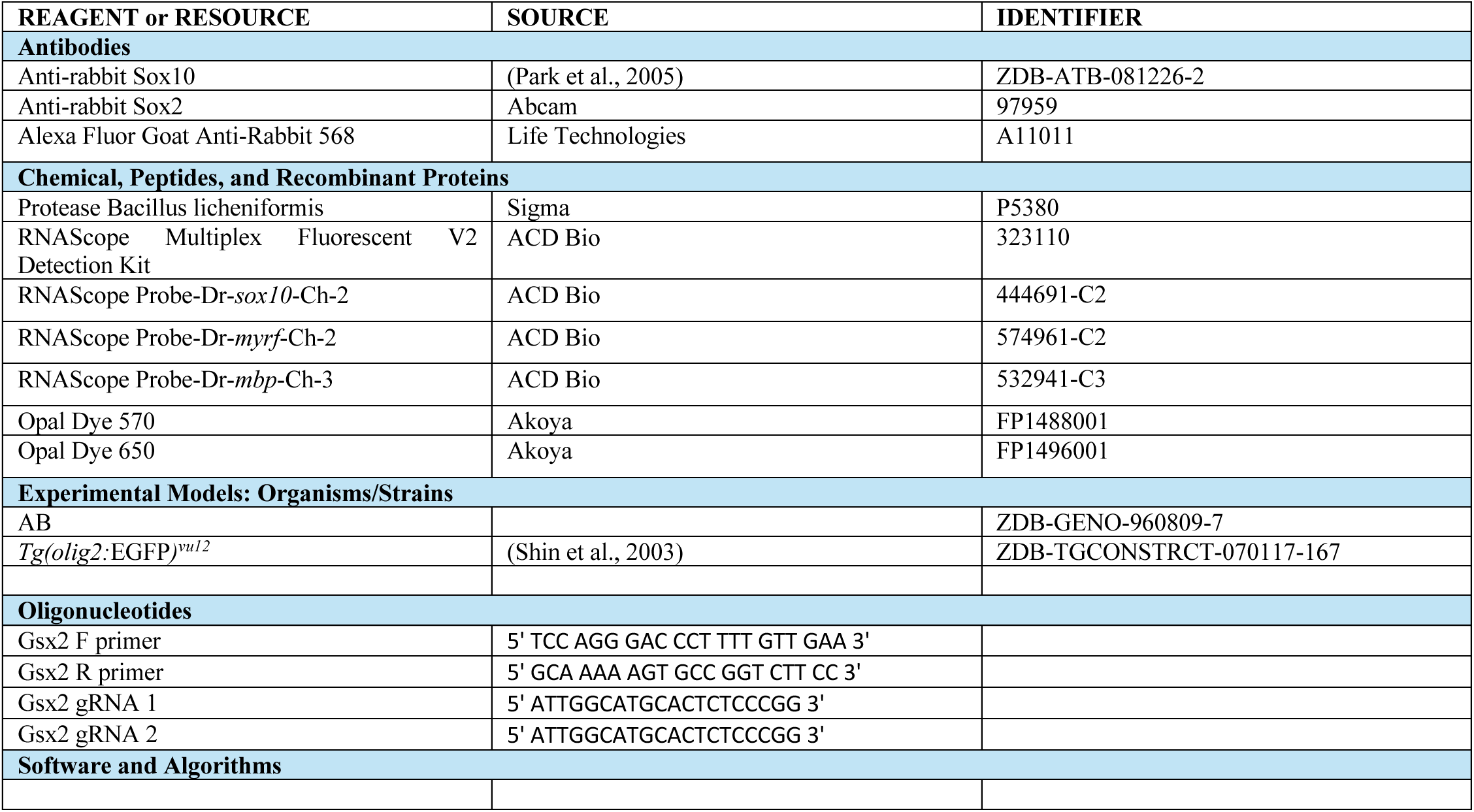

